# Assessing the Intra- and Inter-Subject Reliability of the Perturbational Complexity Index (PCI) of Consciousness for Three Brain Regions Using TMS-EEG

**DOI:** 10.1101/2020.01.08.898775

**Authors:** Kevin A. Caulfield, Matthew T. Savoca, James W. Lopez, Philipp M. Summers, Xingbao Li, Matteo Fecchio, Silvia Casarotto, Marcello Massimini, Mark S. George

## Abstract

**Background:** The perturbational complexity index (PCI) is a useful measure of consciousness that combines transcranial magnetic stimulation (TMS) with electroencephalography (EEG). However, the PCI has not been assessed for reliability between sessions nor is there a clear best stimulation target to acquire a PCI.

**Objective/Hypothesis:** We assessed the reliability of within-subject PCIs between 3 sessions with stimulation over the same premotor, motor, and parietal targets between visits, hypothesizing that we could determine a most reliable TMS-EEG target to acquire PCIs in healthy, conscious adults.

**Methods:** PCIs were acquired for 9 participants (5 women) over 3 sessions within a week. A 64 channel EEG system was used with all electrode impedances ≤ 10kΩ. Premotor, motor, and parietal stimulation targets were identified using a real-time Matlab graphical user interface (GUI). Neuronavigation using an MRI-template brain ensured that every TMS pulse was delivered within 3.0mm and 5° of the stimulation target.

**Results:** Premotor, motor, and parietal PCIs all had significant PCIs (all p < 0.05). However, parietal and motor PCIs had excellent reliability (ICCs = 0.927 and 0.857 respectively) whereas premotor PCIs had good reliability (ICC = 0.737). PCIs were similar between brain sites within each subject in a single visit, but with only a moderate effect (p = 0.024, ICC = 0.480). PCIs on a group level did not differ between brain sites (p = 0.589).

**Conclusions:** The PCI is a reliable measure over this timeframe within each subject for single brain targets. PCIs for parietal and motor sites are most similar between visits. Due to only moderate similarity between PCIs from three brain sites within each session, PCIs should be acquired over at least two brain sites, with parietal and motor regions as the top candidates.

## Introduction

The perturbational complexity index (PCI) is a method of quantifying consciousness by measuring the complexity and integration of brain signals (1). Researchers have defined distinct ranges of the PCI that can be used to differentiate between varying states of consciousness, different stages of sleep, anesthetization, and brain damage (for example, minimally conscious state vs. unresponsive wakefulness)(1-3). PCI values of 0.40 or higher are generally thought to represent consciousness while lower values indicate various states of unconsciousness. While the PCI has made large strides toward the goal of understanding and quantifying consciousness, there are still a few limitations. Currently, the PCI is time consuming and expensive to acquire. PCIs are acquired using transcranial magnetic stimulation (TMS) to perturb the brain and electroencephalography (EEG) to measure the brain’s response. In all studies to date, the PCI has also required each individual’s magnetic resonance imaging (MRI) scan to allow the targeting of specific, individualized gyral targets in real time (1). This set-up is time consuming, bulky, and expensive.

A further area in which PCI can be improved is to rigorously evaluate the reliability of this measure. Researchers have already statistically quantified how the waveforms between TMS-EEG sessions compare, with TMS applied to exactly the same site between visits (confirmed by each subject’s MRI used with neuronavigation)(4). However, the PCI measure itself could use further evaluation to determine if there is consistency between different brain sites within a visit, or between visits within a single brain location.

We had three goals in this study. First, we wanted to determine if a template MRI would be sufficient to use for neuronavigation for use in this TMS-EEG experiment acquiring PCIs.

This could potentially make the TMS-EEG session quicker, easier, and cheaper to implement, allowing it to be more widely accessible. Second, we wanted to assess the reliability of the PCI measure between different brain sites and across 3 visits to determine if there is an optimal stimulation target that yields PCIs that are more reliable between visits. Third, we wanted to determine if there is a minimum number of brain sites that should be stimulated to acquire a representative PCI value, as fewer targets would reduce the overall time of acquisition.

## Materials and Methods

### Study Overview

We enrolled 9 healthy adult participants (5 women, mean age = 26.2, SD = 4.47, range = 22-37) in this three visit IRB-approved study. Each participant gave written, informed consent before starting the experimental protocol. In each visit, participants were set up with an EEG cap and gel before receiving single pulses of TMS at 120% of resting motor threshold (rMT) at three brain locations (premotor, motor, and parietal). The order of stimulation site was randomized and counterbalanced between visits such that each participant received each stimulation site first, second, and third across the three visits. Between visits, the start time of PCI acquisition was within an hour of each session to reduce the potential of circadian differences affecting the level of consciousness between visits. We used the MNI-152 template brain for neuronavigation to ensure that we stimulated precisely the same brain locations with the same coil angle between visits. Detailed procedures for each component of the setup are described in detail below and largely follow previously published methods (1, 2).

### EEG Set-Up

#### EEG Equipment

A 64-channel BrainVision Passive EEG system was used in this experiment (BrainVision, Morrisburg, NC, USA). Two amplifiers connected the EEG cap to a desktop computer using fiber optic cables. These amplifiers were powered with an external battery, and BrainVision Recorder software was used to acquire EEG data at 5000Hz.

#### EEG Cap Placement

We first measured each participant’s head circumference to choose an appropriately sized EEG cap (4 participants were fitted with 56cm caps and 5 were fitted with 58cm caps). Next, we measured nasion-to-inion and tragus-to-tragus lengths and marked their intersection on the scalp. Electrode Cz was placed over the vertex. The EEG cap chinstrap was then tightened to reduce the space between the EEG cap and scalp.

#### Extracephalic EEG Electrode Placement

Three electrodes (reference, EOG1, EOG2) were placed outside of the EEG cap. The reference electrode was fixed to the center of the forehead using double sided EEG electrode tape (Figure 1). EOG1 and EOG2 were affixed above the left and below the right eyes respectively.

**Figure 1:**
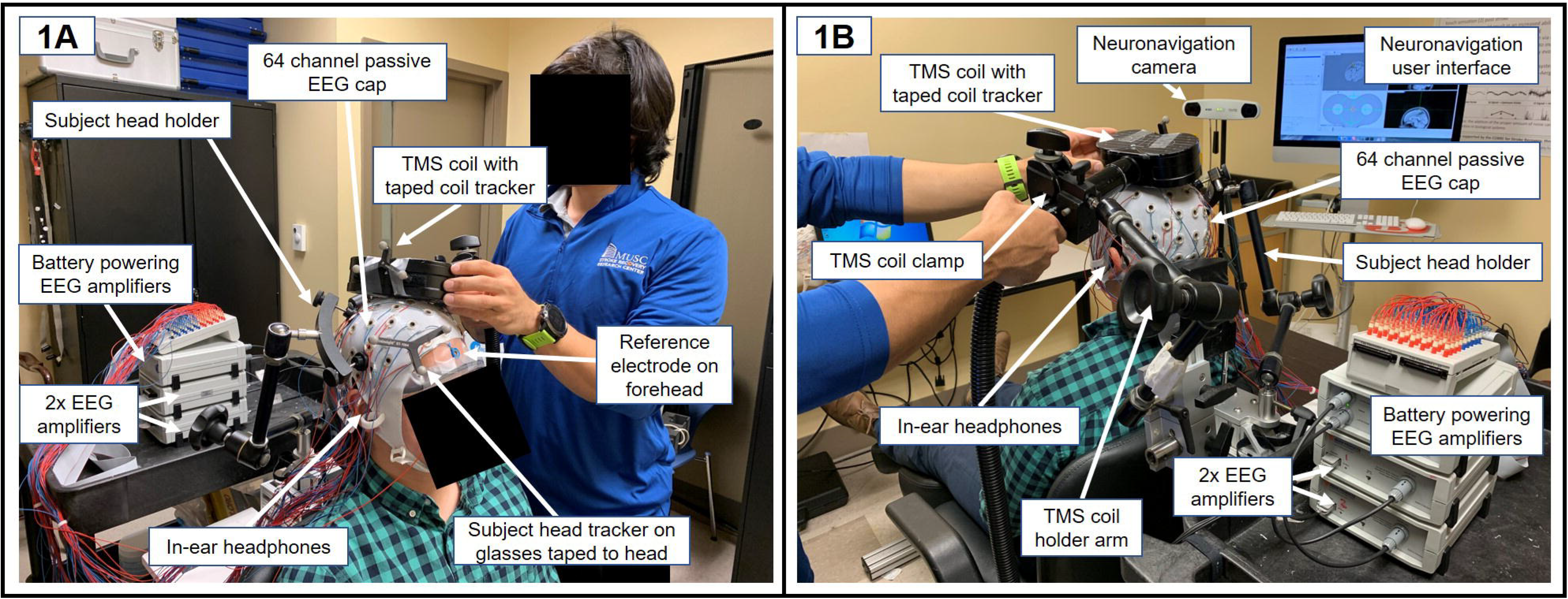
Experimental set-up for TMS-EEG acquisition.

#### EEG Gelling Procedure

Prior to EEG gel being applied to each electrode, we asked participants to brush their hair using a brush we supplied. Two study coordinators gelled the electrodes by first moving hair out of the way using the wooden end of the cotton swab. Once the scalp was visible, the cotton swab was used to apply NuPrep to abrate the scalp. The cotton swab was moved in perpendicular directions in order to not tangle hair or pull hair out of the EEG electrode opening. Following this, study coordinators used a blunt syringe with no tip to insert Electro-Gel into each electrode. Using this protocol, we were able to reduce the impedances of each electrode to ≤ 10kΩ.

### TMS Set-Up

#### Template MRI-Guided Neuronavigation

A goal of this study was to reduce the amount of resources and time it takes to acquire a PCI for healthy, conscious adults. For this reason, we did not MRI scan any participant but rather, stimulation target was informed strictly from brain response using a custom Matlab-based Graphical User Interface (GUI)(2).

Our use of neuronavigation in this study was to mark the coil location and angle to ensure that there was intra- and inter-session reliability between stimulation targets. Using *BrainSight* (Rogue Research, Montreal, Quebec, Canada), we recalibrated the coil prior to each visit and registered each participant’s head in each study visit to the MNI-152 template brain using 5 landmarks: nasion, left/right tragus, and left/right canthus. Each participant wore glasses taped to the cap with an attached subject tracker (**See** Figure 1). We added an additional 9 validation points on each participant’s scalp to control for discrepancies in head shape in each visit, which allows the BrainSight-compatible computer to identify each individual’s scalp. For each participant, every TMS pulse was delivered within 3.0mm and 5° of the target stimulation location.

### TMS Parameters

#### Dosing Procedure: TMS Resting Motor Threshold (rMT)

Using the MagVenture R30 TMS machine with a MagVenture Cool-B65 Butterfly coil (MagVenture, Farum, Denmark), we acquired a resting TMS motor threshold (rMT) through the EEG cap using visual observation for each participant. Once the motor hotspot was located, we titrated the intensity of stimulation delivered through the EEG cap to the amount that elicited 5/10 visual thumb twitches (abductus pollicis brevis muscle). The mean rMT through the EEG cap was 57.89% of machine stimulator output (MSO)(SD = 5.71%, range = 52-71%).

Subsequent stimulation during the experiment was delivered at 120% of each participant’s rMT (mean = 69.33% MSO, SD = 6.81%, range = 62-85%).

#### TMS Stimulation Target Selection

We stimulated over a premotor, motor, and parietal brain target on each of the three days of the experiment. The stimulation target was selected in Visit 1 using established methods that have been previously used for PCI acquisition (1). In line with these previously published methods, we used a real-time Matlab-based Graphical User Interface (GUI) that could visualize the EEG response to single pulses of TMS. Selection of stimulation location was individualized with coil location and angle to elicit the lowest amount of scalp muscle activation with the largest brain response with characteristic frequency of response with each brain area (approximately 25-35Hz for premotor, 15-25Hz for motor, and 5-15Hz for parietal sites)(5, 6). For the premotor location, we stimulated at Brodmann area 6, with the center of the TMS coil placed between electrodes F1, Fz, FC1, and FCz. For motor stimulation we targeted Brodmann area 4, with the center of the coil placed between C1 and Cz. For parietal stimulation, we placed the center of the coil between CP1, CPz, P1, and Pz at Brodmann area 7. Using the Matlab GUI, we changed the coil location in any direction up to a 10mm distance from the start target and allowed an angle change of any percentage that helped to maximize the TMS-EEG signal.

Once an ideal stimulation location was identified, the location and angle of the coil were saved using the BrainSight neuronavigation system. In Visits 2 and 3, each participant’s head was again registered in BrainSight and the same, saved brain locations were targeted in the predetermined counterbalanced order.

#### TMS Stimulation Timing and Number of Pulses

For each brain location, 200 single pulses of TMS were delivered at randomly jittered 0.4-0.5Hz stimulation using a custom Matlab program, replicating previous methods (1). A laptop computer containing the Matlab script was connected directly to the TMS machine using a DAQ in order to externally trigger the TMS pulses. The TMS pulse recharge was offset to 1000ms post-TMS pulse as the window used to calculate PCIs was −600 to 600ms surrounding the pulse at 0ms. TMS pulses were marked with a trigger label “S 1” in the BrainVision Recorder software.

### Noise Masking

As previously reported, noise masking is an essential component for TMS-EEG experiments and failing to adequately mask the noise of the TMS pulse can lead to the presence of an auditory artifact in the EEG signal (7). In this study, we used a scrambled TMS coil noise that constantly plays for noise masking, with the intention of reducing the potential auditory artifact from hearing the TMS coil click. This noise masking paradigm has previously been used to acquire PCIs (1, 2).

The volume of noise masking was individually titrated until each participant reported not being able to hear the TMS coil click at the intensity of 120% rMT, within a safety limit of < 85dB.

## Data Analysis

### EEG Preprocessing

Following TMS-EEG acquisition, the following data processing pipeline was used: First, the datafile was loaded into Matlab (EEGLab) and the ‘S 1’ trigger was used to create individual trials for each TMS pulse, from −600ms before the pulse to 600ms after. The TMS pulse artifact from −2 to 5ms was replaced with baseline EEG signal and a DC correction was run. A first order high pass filter at 1Hz, low pass filter at 80Hz, and notch filter at 59-61Hz was applied to remove low frequency, high frequency, and electrical (60Hz) noise. The data were then down-sampled from 5000Hz to 625Hz. One round of channel and trials rejections preceded a second round of channel rejections before bad channels were interpolated and the reference was re-referenced to the average of all EEG channels. A first round of ICA preceded a second round of trials rejections. Following a second round of ICA, the EEG rater chose which ICA components to reject.

The average number of trials rejections was 27.5 (SD = 18.0, range.= 1-91), average channels rejected were 1.63 (SD = 1.34, range = 0-6)), and average number of ICA rejections was 21.7 (SD = 5.1, range = 8-31).

### PCI Calculation

In an automated data processing step, we used the PCI Lempel-Ziv compression that takes the pre-processed data and outputs a PCI number for each TMS-EEG acquisition session (1). We calculated premotor, motor, and parietal PCIs for each TMS-EEG visit, 3 visits each for 9 participants, for a total of 81 PCI values.

### Statistical Comparisons

We first assessed if there was a group-level difference in PCI score between brain locations or an order effect of stimulation within each visit using two repeated-measures ANOVAs (**See** Figure 2).

**Figure 2:**
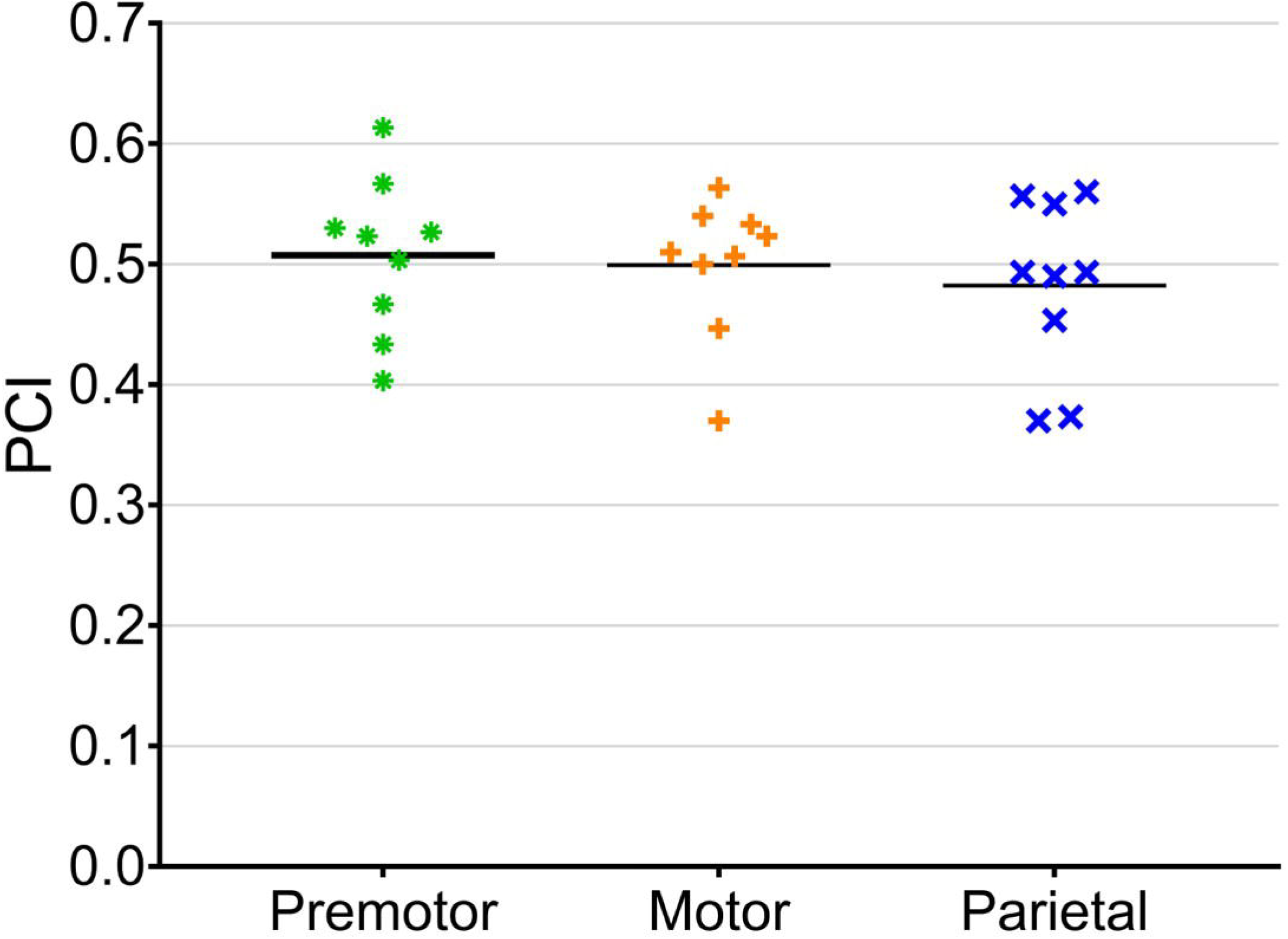
PCIs were not significantly different between brain sites, F(2, 16) = 0.548, p = 0.589, □_p_^2^ = 0.064. Premotor (mean = 0.507, SD = 0.065), motor (mean = 0.499, SD = 0.058), and parietal (mean = 0.482, SD = 0.072) stimulation locations all had similar PCI values.

Next, we used a series of intraclass correlations coefficients (ICCs) to measure how grouped different sets of data were. We ran one ICC for each brain target (premotor, motor, parietal) to assess how similar the PCI values were for each individual between each of the 3 visits (**See** Figure 3). A fourth ICC assessed the similarity level between different brain regions acquired for each participant (**See** Figure 4). A fifth ICC evaluated how grouped the PCIs acquired for premotor, motor, and parietal regions were for each session (**See** Figure 5). All data analyses were conducted in SPSS 25.0.

**Figure 3:**
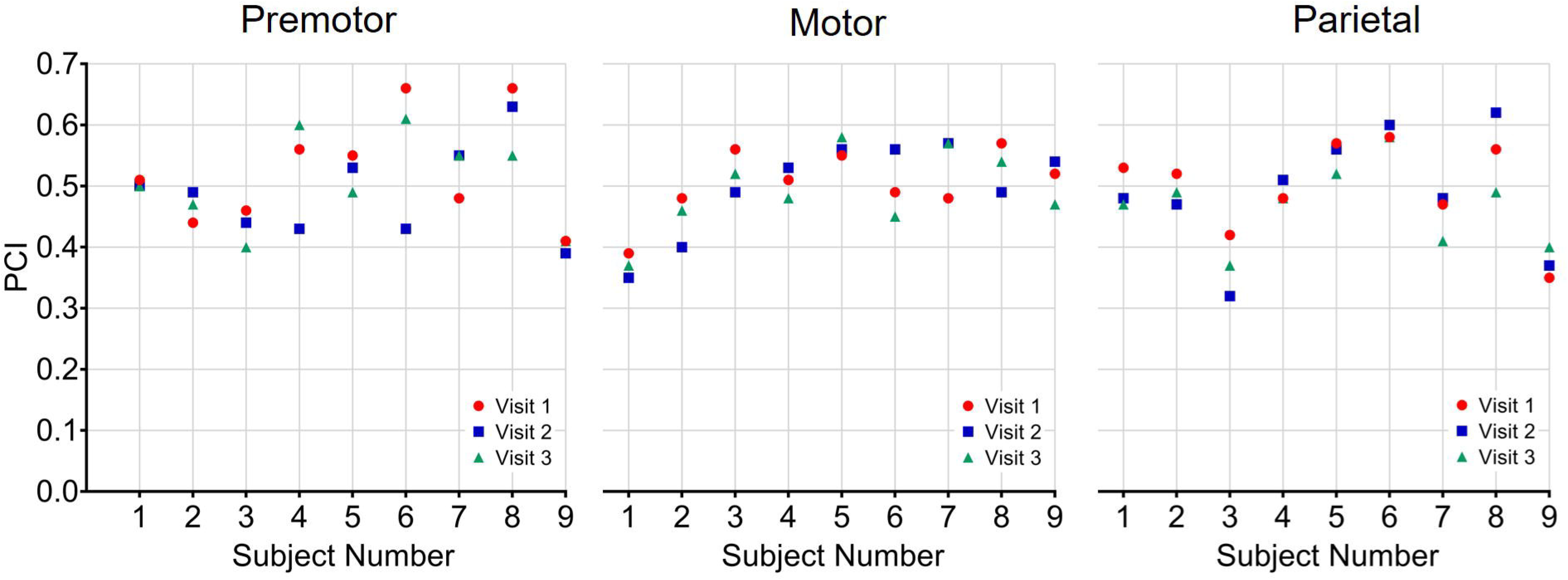
Intraclass correlation coefficients (ICC) for between session PCI reliability were significant for premotor PCIs (ICC = 0.737, F(8,16) = 3.79, p = 0.011), motor PCIs (ICC = 0.857, F(8,16) = 6.45, p = 0.001), and parietal PCIs (ICC = 0.927, F(8,16) = 14.83, p < 0.001).

**Figure 4:**
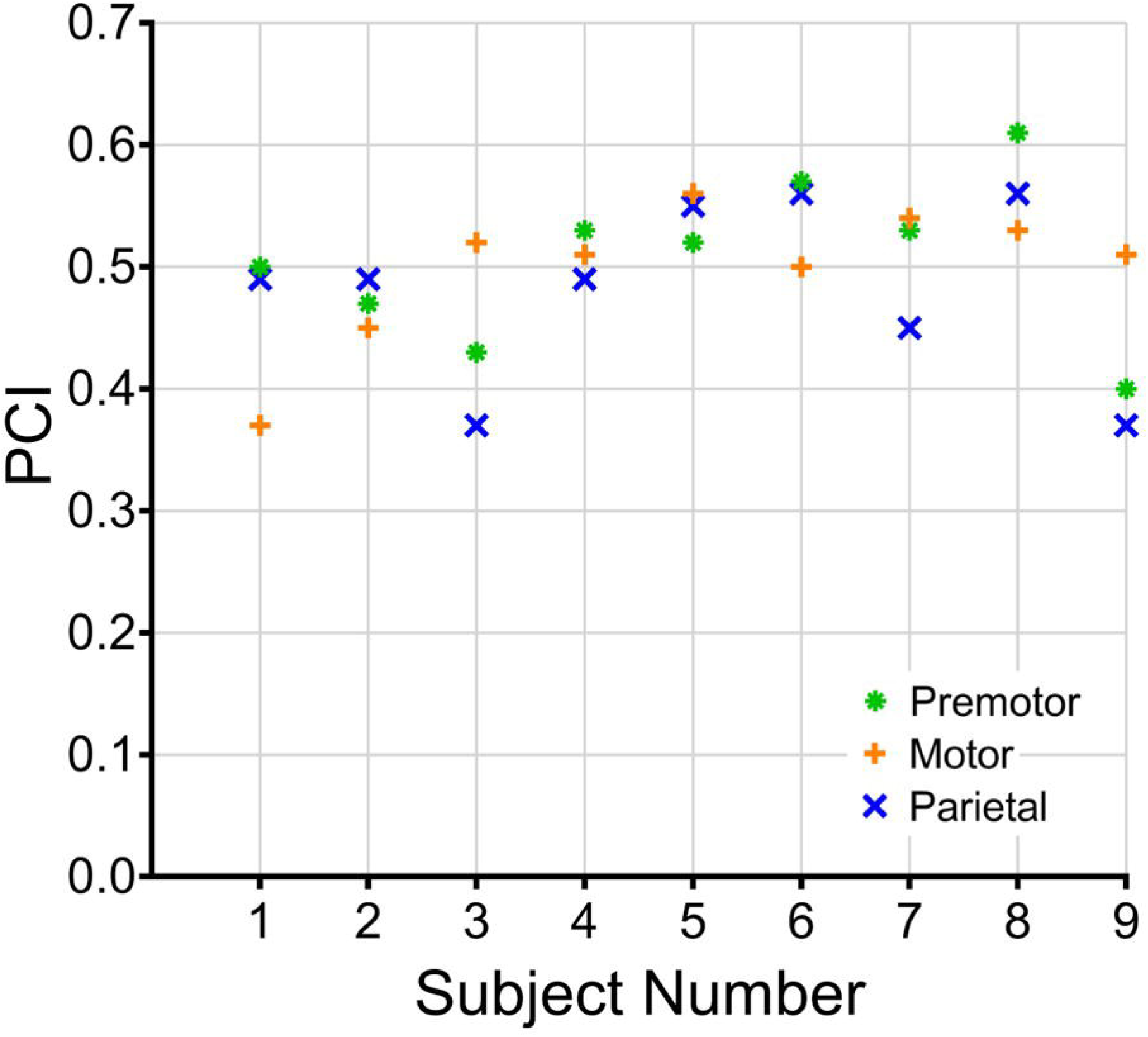
The averaged PCI values between brain sites for each individual were moderately similar, ICC = 0.646, F(8, 16) = 2.74, p = 0.041.

**Figure 5:**
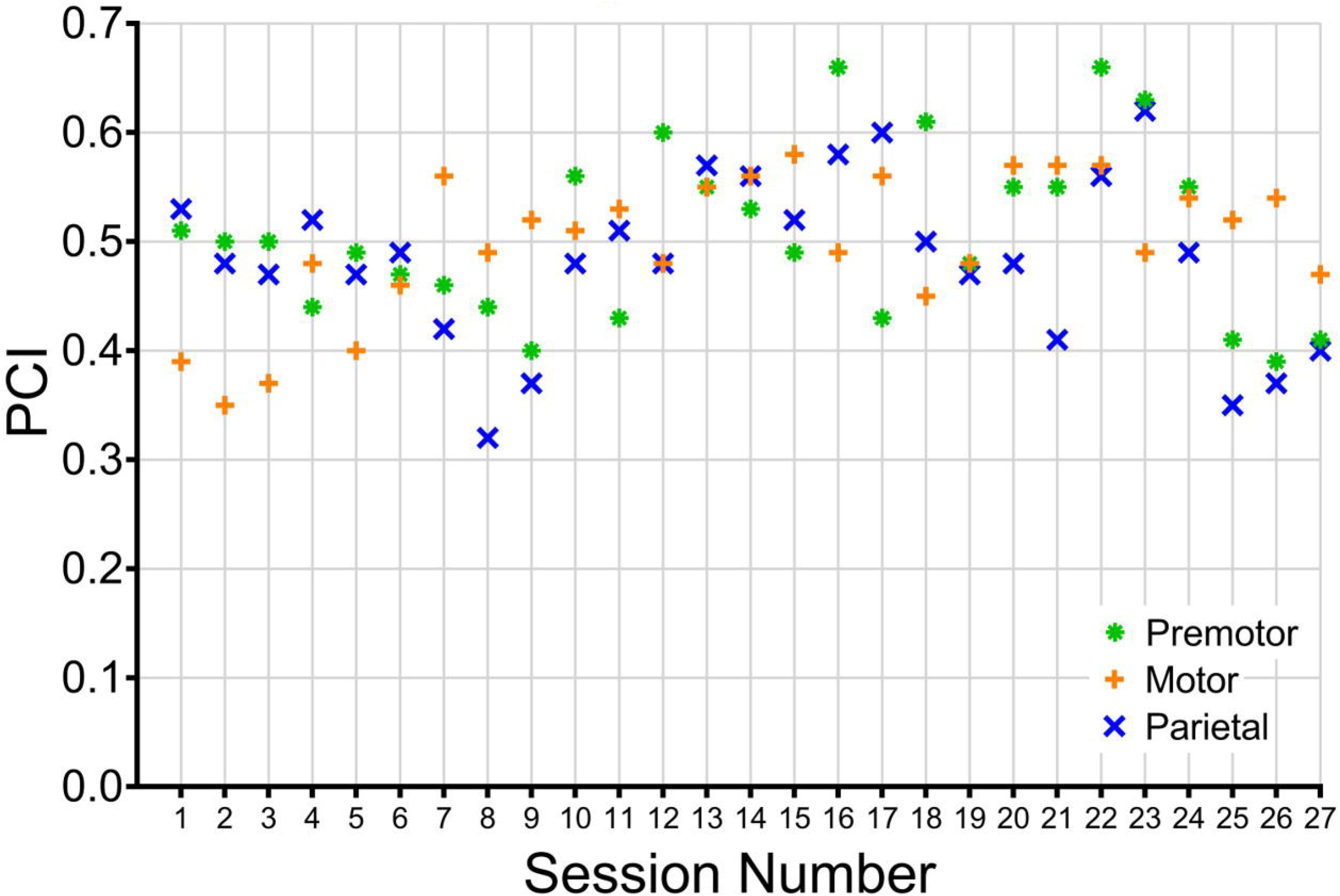
The PCI values for different brain sites within a single visit was significant, but with only poor/moderate similarity between brain sites, ICC = 0.480, F(26, 52) = 1.90, p = 0.024.

## Results

### ANOVA: Order Effects

There were no order effects of stimulation, F(2, 25) = 0.375, p = 0.689, □_p_^2^ = 0.029. The PCI values for the first (mean = 0.490, SD = 0.063), second (mean = 0.505, SD = 0.074), and third (mean = 0.494, SD = 0.084) sites stimulated were non-statistically different.

### ANOVA: Do PCIs Differ Between Brain Sites on a Group Level?

PCIs did not significantly differ between brain sites, F(2, 25) = 1.72, p = 0.199, □_p_^2^ = 0.121. Premotor (mean = 0.507, SD = 0.065), motor (mean = 0.499, SD = 0.058), and parietal (mean = 0.482, SD = 0.072) stimulation locations all had similar PCI values (**See** Figure 2).

### ICCs: Is There an Optimal Stimulation Target for Acquiring Reliable PCIs? Assessing the Similarity of PCIs Within a Brain Region, Within-Subject, Between Visits 1-3

We used three ICCs to evaluate the reliability of PCI within each individual, at each brain location, between visits. Each ICC was significant. Parietal PCIs had the highest ICC; Parietal ICC = 0.927, F(8,16) = 14.83, p < 0.001 (**See** Figure 3). The next most significant ICC was motor, Motor ICC = 0.857, F(8,16) = 6.45, p = 0.001 (**See** Figure 3). Lastly, premotor PCIs had lowest reliability, Premotor ICC = 0.737, F(8,16) = 3.79, p = 0.011 (**See** Figure 3). While each ICC was significant, there was stratification in the reliability of ICC; Parietal and motor ICCs had “excellent reliability,” while Premotor ICC had “good reliability” (8).

### ICC: How Similar Are Averaged PCI Values Within Each Subject, Between Brain Sites?

A key question when considering the time of TMS-EEG acquisition and PCI calculation is how many brain sites need to be stimulated to acquire a reliable PCI. Thus, we used an ICC to assess how similar the average PCI values at each brain region was to each other for each participant. There was significant, moderate reliability between brain sites for each individual, ICC = 0.646, F(8, 16) = 2.74, p = 0.041 (**See** Figure 4). This result suggests that the PCI values within each individual and between brain sites are somewhat similar, with good reliability (8).

### ICC: How Similar Were PCIs Within Each Session, Between Brain Sites?

Lastly, it is important to determine the similarity of PCIs between brain sites, within a single day’s visit to inform how many stimulation sites should be targeted in each session to acquire a PCI. To measure this, we conducted an ICC assessing the reliability of PCIs within each session between visits (N = 27). This ICC was significant, ICC = 0.480, F(26, 52) = 1.90, p = 0.024 (**See** Figure 5). However, the ICC had only fair reliability (8).

## Discussion

In this study, we assessed the reliability of PCIs in 9 healthy, conscious adults, over 3 visits, in premotor, motor, and parietal locations in each visit. We were able to calculate a PCI value for each of these sessions and were able to assess the similarity of PCIs in different settings. Our method of using BrainSight neuronavigation using a template MRI enabled us to ensure that stimulation targeted exactly the same spot within and between visits without using each participant’s structural MRI scan. This reduces the cost of acquiring PCIs and may make the use of PCIs more widely accessible.

PCIs for parietal and premotor sites were the most reliable for within-individual, between visit PCIs, and PCIs did somewhat differ between brain sites within individual visits, showing only moderate amounts of similarity with ICC analyses. Taken in sum, these data suggest that at least two brain sites should be targeted within a single day to acquire a representative PCI value for an individual. If there is a choice of where to stimulate (i.e. if there is no brain damage limiting the stimulation target to intact cortex), it appears best to use TMS over parietal and motor regions as these data were most reliable. This recommendation may be particularly useful for investigating healthy adults in different states of consciousness or in neurological or psychiatric conditions that are not associated with damaged cortex.

There were several limitations of this study. While taking multiple PCI measurements likely increases the chances of obtaining a ground truth PCI value, there was no way to assess how different the PCI at each brain site was from each participant’s true PCI. Thus, while we can assess the reliability of PCI values at and between each brain site, it is not possible to compare these to a true PCI. It also continues to be unclear whether it is the PCI that differs by brain site, or if it is the measurement of PCI that differs by brain site.

There have also been some recent methodological developments that could potentially improve the TMS-EEG signal-to-noise ratio that we either chose not to implement or were developed after we vegan the study. For example, some researchers now individually titrate the noise masking frequency to more fully mask the noise of the TMS click using custom Matlab scripts that did not previously exist but may potentially reduce the auditory artifact that can be present in TMS-EEG recordings (**unpublished work**). This potentially reduces the auditory artifact that can be present in TMS-EEG recordings. In addition, some neuronavigation systems are able to prevent the TMS coil from delivering a pulse if it is not within a certain distance or angle of the target. While we ensured that stimulation was always delivered within 3.0mm and ≤ 5° of the stimulation target, it could be possible to more rigorously control for the stimulation target and angle based on the neuronavigation system used.

A further limitation was our lack of use of a questionnaire that may have been able to link small changes in healthy, conscious PCIs with micro-changes in brain states between TMS-EEG acquisitions, both within and between visits. We chose to not implement a questionnaire as it was not the primary focus of this study. However, it would have been useful to confirm how conscious each participant was before each TMS-EEG acquisition and could be worth including in future experiments.

Future methodological work should also consider studying the difference between PCI values at the same brain site multiple times within a single visit. This could further refine the sensitivity of PCI values that would not be predicted to change much on a micro-, minute-to-minute basis. Lastly, it would be useful to methodically quantify how stimulation at higher doses (e.g. 140% or 160% of rMT) would impact the brain response to TMS and the PCI, especially between different states of consciousness.

## Conclusions

The PCI is a reliable measure in conscious, healthy adults, particularly at parietal and motor brain targets. Between visits, the PCI values for parietal, motor, and premotor locations were significantly reliable within individuals. We demonstrated that the PCI can be acquired using a template-brain for neuronavigation to ensure that stimulation is delivered at exactly the same location in each visit. Stimulating over at least two brain sites appears to be the best practice for acquiring PCIs, as even within a single visit, there can be a range of PCI values within an individual depending on the location stimulated.

## References

1. Casali AG, Gosseries O, Rosanova M, Boly M, Sarasso S, Casali KR, et al. A theoretically based index of consciousness independent of sensory processing and behavior. Science translational medicine. 2013;5(198):198ra05.

2. Casarotto S, Comanducci A, Rosanova M, Sarasso S, Fecchio M, Napolitani M, et al. Stratification of unresponsive patients by an independently validated index of brain complexity. Annals of neurology. 2016;80(5):718–29.

3. Sarasso S, Boly M, Napolitani M, Gosseries O, Charland-Verville V, Casarotto S, et al. Consciousness and Complexity during Unresponsiveness Induced by Propofol, Xenon, and Ketamine. Current biology: CB. 2015;25(23):3099–105.

4. Casarotto S, Romero Lauro LJ, Bellina V, Casali AG, Rosanova M, Pigorini A, et al. EEG responses to TMS are sensitive to changes in the perturbation parameters and repeatable over time. PLoS One. 2010;5(4):e10281.

5. Fecchio M, Pigorini A, Comanducci A, Sarasso S, Casarotto S, Premoli I, et al. The spectral features of EEG responses to transcranial magnetic stimulation of the primary motor cortex depend on the amplitude of the motor evoked potentials. PLoS One. 2017;12(9):e0184910.

6. Bodart O, Fecchio M, Massimini M, Wannez S, Virgillito A, Casarotto S, et al. Meditation-induced modulation of brain response to transcranial magnetic stimulation. Brain Stimul. 2018;11(6):1397–400.

7. Conde V, Tomasevic L, Akopian I, Stanek K, Saturnino GB, Thielscher A, et al. The non-transcranial TMS-evoked potential is an inherent source of ambiguity in TMS-EEG studies. Neuroimage. 2019;185:300–12.

8. Cicchetti D. Guidelines, Criteria, and Rules of Thumb for Evaluating Normed and Standardized Assessment Instrument in Psychology. Psychological Assessment. 1994;6:284–90.

